# Chemical proteomics suggests sofosbuvir is metabolically attached to proteins through a UMPylation-like pathway

**DOI:** 10.64898/2026.10.17.759838

**Authors:** Manuel Hertwig, Pavel Kielkowski

**Affiliations:** LMU Munich, Department of Chemistry, Butenandtstr. 5-13, 81377 Munich, Germany

**Keywords:** Sofosbuvir, phosphoramidate prodrugs, Post-translational modifications, chemical proteomics, UMPylation

## Abstract

Sofosbuvir is a pronucleotide prodrug (ProTide) that releases a uridine monophosphate analogue, which could modify proteins chemically or through UMP-transferring enzymes. Using an alkyne-tagged probe and chemical proteomics, we identified about 120 labelled proteins and a UMPylation-like (+774.31 Da) modification on cathepsin Z, suggesting that nucleotide prodrugs may enter protein nucleotidylation pathways.

## Introduction

Sofosbuvir (**Figure 1A**) is a nucleotide analogue prodrug that has transformed the treatment of chronic hepatitis C virus (HCV) infection, with cure rates above 90% across genotypes^1^. It acts as a chain terminator of transcription by inhibiting the viral RNA-dependent RNA polymerase NS5B after intracellular conversion to its active triphosphate form, 2’-deoxy-2’-fluoro-2’-C-methyluridine triphosphate^2,3^. The parent nucleoside is a poor substrate for cellular kinases, so the initial phosphorylation step is inefficient and limits the formation of active metabolites. Sofosbuvir overcomes this limitation through the ProTide approach^4,5^, in which the monophosphate is masked as a phosphoramidate carrying an aryl group and an amino acid ester. This masking neutralizes the charge of the phosphate, thereby enabling efficient cell entry, and bypasses the first phosphorylation step. The ProTide strategy has since been applied well beyond HCV^4^. Tenofovir alafenamide, a phosphonamidate prodrug of the acyclic nucleotide analogue tenofovir, is used against HIV and hepatitis B virus^6^. Remdesivir, a phosphoramidate prodrug of a 1’-cyano-substituted C-adenosine analogue, was developed against Ebola virus and became a widely used antiviral against SARS-CoV-2^7,8^. Further examples include bemnifosbuvir (AT-527), a guanosine analogue prodrug evaluated against HCV and SARS-CoV-2, and, outside virology, the gemcitabine and 5-fluoro-2’-deoxyuridine ProTides NUC-1031 and NUC-3373, which have been investigated as anticancer agents^9,10^. Inside the cell, and predominantly in hepatocytes, the prodrug is unmasked in a stepwise process involving carboxylesterase 1 or cathepsin A^11^, spontaneous cyclization with loss of the aryl group^12^, and cleavage of the P-N bond by phosphoramidases, which release the uridine monophosphate analogue (**Figure 1B**) ^4^. Subsequent phosphorylation by UMP-CMP kinase and nucleoside diphosphate kinase yields the active triphosphate. Because this pathway generates a UMP analogue inside the cell, we asked whether it could also be used by enzymes that transfer UMP onto proteins, and whether sofosbuvir-derived metabolites might therefore modify proteins in a manner resembling UMPylation^13^. Protein AMPylation is a post-translational modification in which AMP from ATP is covalently attached to hydroxyl-containing amino acid side chains, mainly serine, threonine and tyrosine, and it is catalysed by Fic-domain enzymes and the mitochondrial pseudokinase protein adenylyltransferase SelO (SELENOO)^14–16^. In human cells, the best-characterised AMPylator FICD regulates the unfolded protein response by modifying the ER chaperone BiP (HSPA5)^17,18^, and AMPylation has also been linked to neuronal development and neurodegeneration^19–22^. The related UMPylation, the transfer of UMP from UTP, is much less understood and has so far been described for only a few proteins, such as the bacterial PII protein GlnK, and *in vitro* for some AMP-transferases that accepts UTP as a substrate^23–26^. To study AMPylation in living cells, we previously developed proN6pA, an N6-propargyl adenosine phosphoramidate probe^19,27,28^. Like sofosbuvir, it is a ProTide, which delivers the nucleoside monophosphate analogue into the cell and bypasses the first phosphorylation step. After intracellular activation and conversion to the ATP analogue, it is transferred onto proteins by AMPylating enzymes, and the terminal alkyne enables enrichment and identification of the modified proteins. Following the same principle, a uridine-based phosphoramidate probe, proN3pU, was developed to profile protein UMPylation, and it enabled the identification of proteins modified by UMP in living cells^24^. Because the intracellular activation of sofosbuvir also generates a UMP analogue, we asked whether sofosbuvir-derived metabolites might be used by enzymes that transfer nucleotides onto proteins, and thus modify proteins in a manner resembling UMPylation. In parallel to this productive pathway, the cyclic intermediate is activated towards nucleophilic attack and may react with nucleophilic amino acid residues of cellular proteins, which in principle could result in nonspecific covalent protein labelling ^12^. Although sofosbuvir primarily inhibits the viral RNA polymerase NS5B, it may therefore also interact with human proteins. Such off-targets can be deconvoluted by activity-based protein profiling combined with LC-MS/MS chemical proteomics, which enables an unbiased, proteome-wide characterisation of interactions between bioactive small molecules and proteins^29,30^. This approach relies on a probe, a derivative of the small molecule equipped with a terminal alkyne. Proteins that form a covalent bond with the probe are conjugated to a biotin tag by Cu(I)-catalysed azide-alkyne cycloaddition (CuAAC)^31,32^ and enriched on streptavidin-coated paramagnetic beads^33^. Tryptic digestion of the enriched proteins and subsequent LC-MS/MS analysis then identify the proteins covalently modified by the probe. Direct identification of sofosbuvir-protein adducts by LC-MS/MS proteomics is, however, practically unfeasible: the adducts are of very low abundance, the phosphate group ionizes poorly, and the N-glycosidic (C-N) bond fragments readily during MS/MS, which hampers both detection and localisation of the modification site^34,35^. Enrichment-based chemical proteomics therefore offers an indirect route to the modified proteins. In this study, we performed chemical proteomics profiling of sofosbuvir in living cells using the alkyne-tagged probe N3pSof.

**Figure 1.**
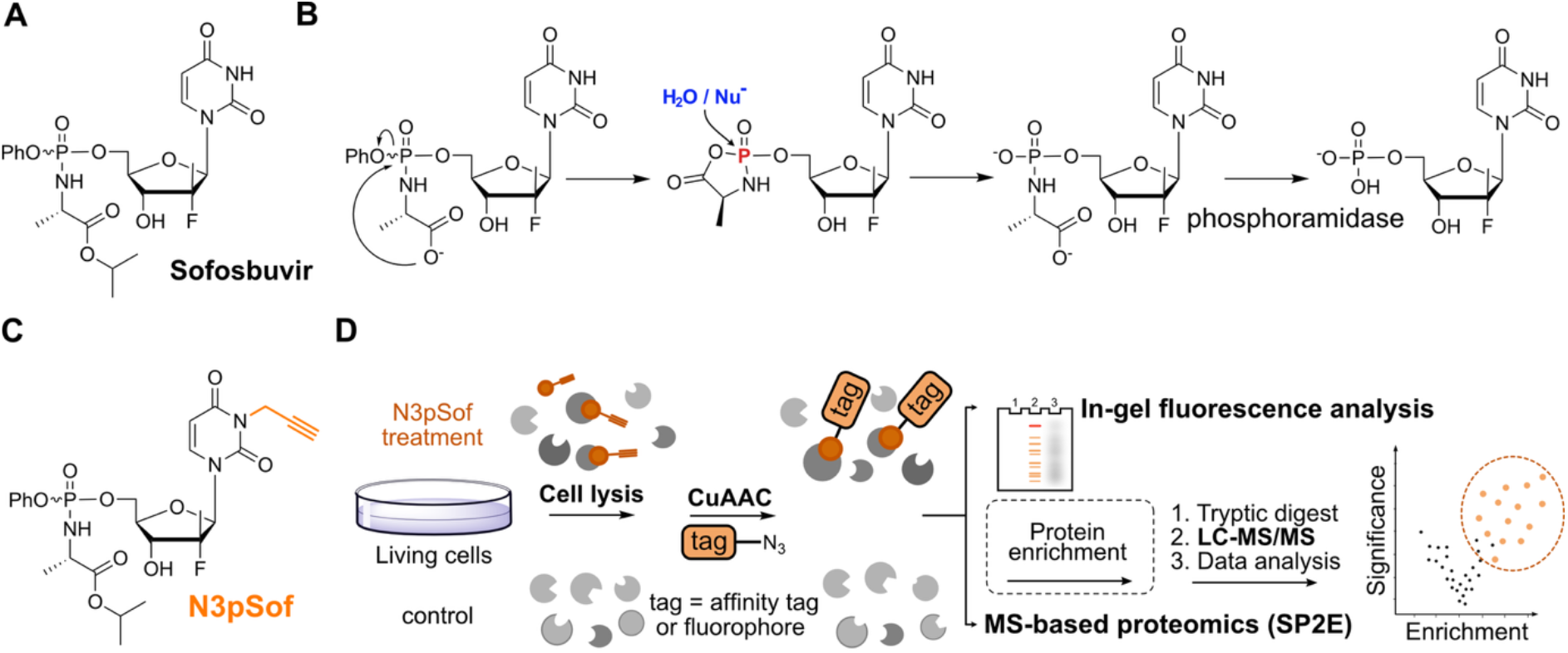
**A**) Sofosbuvir structure. **B**) ProTides activation mechanism. **C**) Structure of the N3pSof probe. **D**) General scheme of chemical proteomics workflow.

## Results

We have designed and synthetized the *N*^3^-propargyl sofosbuvir probe (N3pSof, **Figure 1C**). The synthesis was done analogously to previous preparation of a uridine prodrug probe for UMPylation profiling by a one step alkylation reaction (Figure S1). Next, we continued with the in-gel fluorescence analysis and mass spectrometry-based proteomics of N3pSof enriched proteins (**Figure 1D**). HEK293T cells were treated with increasing concentrations of N3pSof, and the lysates were subjected to CuAAC with TAMRA-azide, followed by SDS-PAGE. Protein labelling was detectable at 50 µM and increased to 200 µM (**Figure 2A**). The labelling was, however, weaker than that of the proN6pA probe, which is used to study protein AMPylation (**Figure 2A**). To identify the labelled proteins, we performed LC-MS/MS chemical proteomics using the solid-phase-enhanced sample preparation for enrichment (SP2E) workflow^33^. In brief, the lysates of probe-treated cells are conjugated to biotin-azide by CuAAC, and the proteins are then captured on paramagnetic carboxylate-coated beads and purified by organic solvent-driven precipitation, which removes excess click reagents and other contaminants. The biotinylated proteins are subsequently enriched on streptavidin beads, washed under stringent conditions, and digested with trypsin, and the resulting peptides are analysed by LC-MS/MS. We applied this workflow to HEK293T cells treated with 100 µM or 200 µM N3pSof for 16 h, using four biological replicates per condition. The data-independent acquisition data were processed with DIA-NN^36^ and analysed with limma and limpa^37,38^ using a custom R script. Overall, 20 and 51 proteins were significantly enriched by 100 µM and 200 µM N3pSof, respectively (**Figure 2B** and S2). The most significantly enriched proteins included prostaglandin E synthase 2 (PTGES2), cathepsin B (CTSB), monoacylglycerol lipase ABHD6 (ABHD6), N-sulfoglucosamine sulfohydrolase (SGSH), phospholipase D family member 3 (PLD3), and protein phosphatase methylesterase 1 (PPME1).

**Figure 2.**
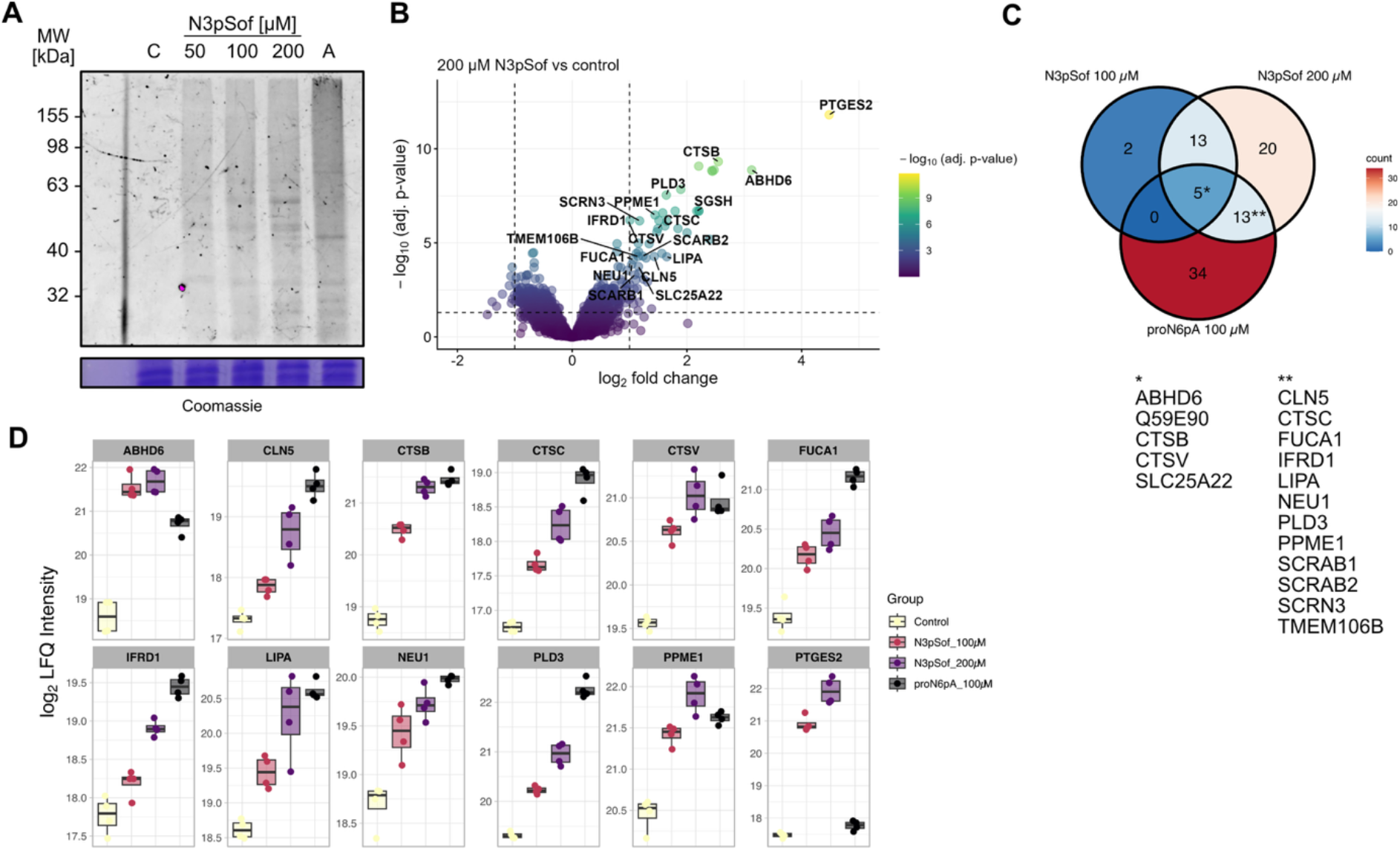
**A**) In-gel fluorescence analysis using the N3pSof or pro-N6pA (A, 100 μM) probe; C – control (cells treated with plain DMSO). **B**) Volcano plot visualizing the proteins enriched by 200 µM N3pSof probe (*n* = 4 biological replicates, cut-off lines: adj. *p*-value = 1.3, fold change = 1). **C**) Venn diagram showing the overlap of significantly enriched proteins between 100 and 200 μM N3pSof and 100 μM pro-N6pA probe.

To exclude that the enrichment merely reflected changes in protein abundance, we analysed the whole proteome of cells treated with sofosbuvir or N3pSof using single-pot solid-phase-enhanced sample preparation (SP3) ^39,40^. No significant changes in protein levels were observed upon either treatment (Figure S3 and S4), including for the enriched proteins, which indicates that the enrichment results from probe labelling and not from altered expression. To place these results in the context of protein nucleotidylation, we performed a back-to-back comparison with cells treated with 100 µM proN6pA (Figure S5). Notably, several proteins were enriched by all three treatments (100 and 200 µM N3pSof and 100 µM proN6pA, **Figure 2C**), which suggests that the uridine- and adenosine-based ProTide probes target a common set of proteins. Profile plots of the individual proteins showed that N3pSof enriched them in a concentration-dependent manner (**Figure 2D**), as expected for a probe-dependent covalent modification. For the proteins shared with proN6pA, however, the enrichment by N3pSof did not reach the efficiency of the proN6pA probe. With the exception for PTGES2, which was only enriched by N3pSof (**Figure 2D**). This overlap might be consistent with the hypothesis that sofosbuvir-derived metabolites enter a protein modification pathway that resembles AMPylation and, by extension, UMPylation. Taken together, the in-gel fluorescence and chemical proteomics data show that N3pSof labels a defined set of proteins in a concentration-dependent manner, and that part of this set overlaps with the targets of the AMPylation probe proN6pA, consistent with a putatively shared nucleotidylation-like pathway.

To validate cathepsin B (CTSB) as a target, we transiently overexpressed wild-type CTSB or CTSB point mutants in HEK293T cells, and repeated the SP2E enrichment. This time, we eluted the captured proteins from the streptavidin-coated beads, and analysed the eluates by in-gel fluorescence analysis and subsequent western blot (**Figure 3A** and **B**). The mutants comprised a substitution of the active-site cysteine (C108A) and substitutions of two serine residues (S104A and S107G) that had previously been reported to be AMPylated^19^. A strong fluorescent band at the molecular weight of CTSB was observed for wild-type CTSB, whereas no such band was detected for either of the mutants (**Figure 3A**). Western blot analysis of non-enriched whole lysates showed comparable expression of the wild-type and mutant proteins, and both the full-length and the processed forms of CTSB were present in all cases, indicating that the mutations did not impair expression or proteolytic processing (Figure S6). The loss of labelling in the mutants therefore indicates that these residues are required for the modification, although we cannot fully exclude that the C108A substitution affects the folding or stability of CTSB. To confirm the identity of the labelled band, we conducted a western blot with an anti-CTSB antibody (**Figure 3B**). Again, a strong CTSB signal for both the single chain protein (∼ 33 kDa) and the heavy chain (∼ 28 kDa)^41^ was observed in the cells overexpressing wild-type CTSB, but not in the cells expressing the mutants.

**Figure 3.**
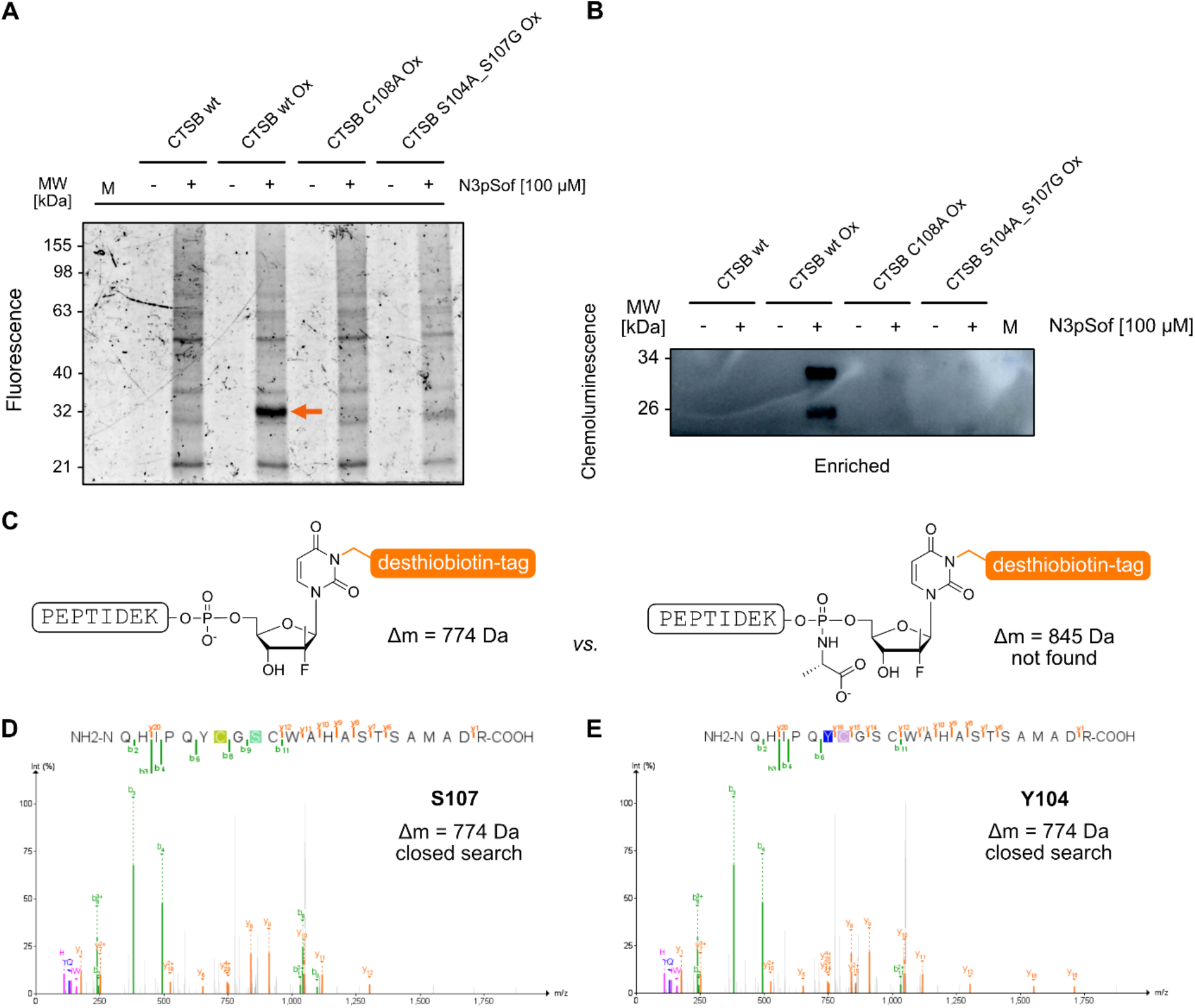
**A**) In-gel fluorescence analysis of N3pSof treated and enriched HEK293T cell lysates containing endogenous CSTB only, transiently overexpressed wild-type CTSB and CTSB point mutants. **B**) Western blot analysis of N3pSof labelling in HEK293T cells from (A) after the SP2E pull-down. **C**) Two anticipated structures of modified peptides resulting from metabolic labelling (left, Δm = 774 Da) and chemical labelling (right, Δm = 845 Da). **D**) and **E**) Selected MS^2^ spectra of CTSZ active site with N3pSof modified peptide on serine 107 and tyrosine 104.

We next attempted to identify the modification sites on CTSB. Immunoprecipitation of overexpressed CTSB from N3pSof-treated cells followed by LC-MS/MS analysis did not yield any modified peptides. We therefore employed a site-identification (site-ID) workflow^42,43^ using desthiobiotin-azide instead of biotin-azide. The principle is that desthiobiotin binds streptavidin with a markedly lower affinity than biotin, so desthiobiotin-modified peptides can be released under mild conditions, whereas biotinylated peptides could not be eluted efficiently. The probe-labelled proteins were conjugated to desthiobiotin-azide by CuAAC, enriched on streptavidin beads, and digested on-bead with trypsin. The unmodified tryptic peptides were removed by stringent washing, and only the probe-modified, desthiobiotin-tagged peptides were subsequently released from the beads with 50% and 80% acetonitrile, respectively. The eluted peptides were analysed by data-dependent acquisition LC-MS/MS, with MS1 and MS2 spectra recorded at high resolution in the Orbitrap. The data were searched with the closed search of the MSFragger suite^44,45^, using variable mass shifts corresponding to the expected modifications. For the metabolic labelling, resembling UMPylation, the calculated mass shift was +774.31 Da. For the proposed chemical labelling, in which the cyclic intermediate of the ProTide activation is opened by a nucleophilic amino acid residue of a protein, it was +845.35 Da (**Figure 3C**). Both mass shifts include the desthiobiotin linker, which remains attached to the modified peptide. The two shifts differ by 71.04 Da, the mass of an alanine residue, which reflects the alanine of the ProTide being retained in the ring-opened adduct but not in the UMP-like product. We searched for both modifications on serine, tyrosine and other nucleophilic residues, including cysteine. This analysis identified modification of serine 107 (S107) and tyrosine 104 (Y104) of cathepsin Z (CTSZ) (**Figure 3D** and **E**), both with a mass shift of +774.31 Da, consistent with a UMPylation-like modification rather than chemical labelling by the ring-opened intermediate. No modified cysteine was detected. Taken together, these results indicate that sofosbuvir-derived metabolites modify cysteine cathepsins in a UMPylation-like manner rather than through nonspecific chemical labelling.

## Discussion

In this study, we show that the sofosbuvir probe N3pSof covalently attaches to a set of proteins in living cells, and that the modification of CTSZ at S107 and Y104 corresponds to a UMPylation-like adduct. Comparison of the N3pSof-enriched proteins with the published AMPylation and UMPylation probe datasets revealed an overlap of three proteins (ABHD6, CTSB and CTSV) between all four datasets and of 18 proteins including PTGES2, AIG1 and CES2 between N3pSof and the UMPylation probe (Figure S7). The larger overlap with the UMPylation dataset is in line with the uridine nature of the sofosbuvir-derived metabolite. We propose that N3pSof labels proteins through two possible routes: metabolic incorporation of the activated UMP analogue by nucleotidylating enzymes^46^, and chemical labelling by the reactive cyclic intermediate of the ProTide activation, which is opened by nucleophilic residues of proteins (Figure S7). In conclusion, we show that the ProTide drug sofosbuvir can label proteins in living cells, and that it is attached to cathepsin Z in a manner consistent with a UMPylation-like modification. Our findings suggest that nucleotide analogue prodrugs may enter protein nucleotidylation pathways in addition to their intended antiviral mechanism, and they provide a basis for investigating the enzymes involved and the functional consequences of this modification.

## Acknowledgements

We are grateful for the support from the Deutsche Forschungsgemeinschaft (DFG, German Research Foundation) with funds from SFB1309 (Chemical Biology of Epigenetic Modifications), project 325871075 (P. K.); P.K. were further supported by the BMFTR in the framework of the Cluster4Future program (Cluster for Nucleic Acid Therapeutics Munich, CNATM) (Project ID: 03ZU1201AA) and Boehringer Ingelheim Foundation – Plus 3 Program.

## Data availability

Supporting Information (PDF) contains supporting figures, detail description of the organic synthesis, biochemical methods, proteomic methods and list of reagents used in the study.

All MS data are available via ProteomeXchange with identifier PXD085731.

Token: KNuE2iCMgWl8

