## Supporting Information for "Chemical proteomics suggests sofosbuvir is metabolically attached to proteins through a UMPylation-like pathway"

#### Supplementary Figures

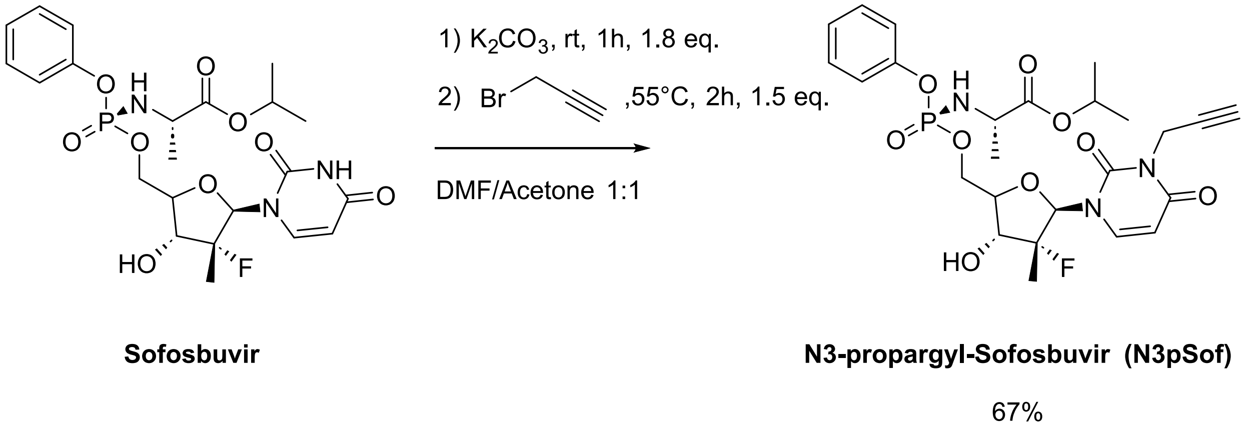

**Figure S1. Synthesis of the N3pSof probe.** N3pSof was obtained from commercially available Sofosbuvir in a one-step synthesis involving deprotonation followed by an $S_n2$ reaction at the imide nitrogen.

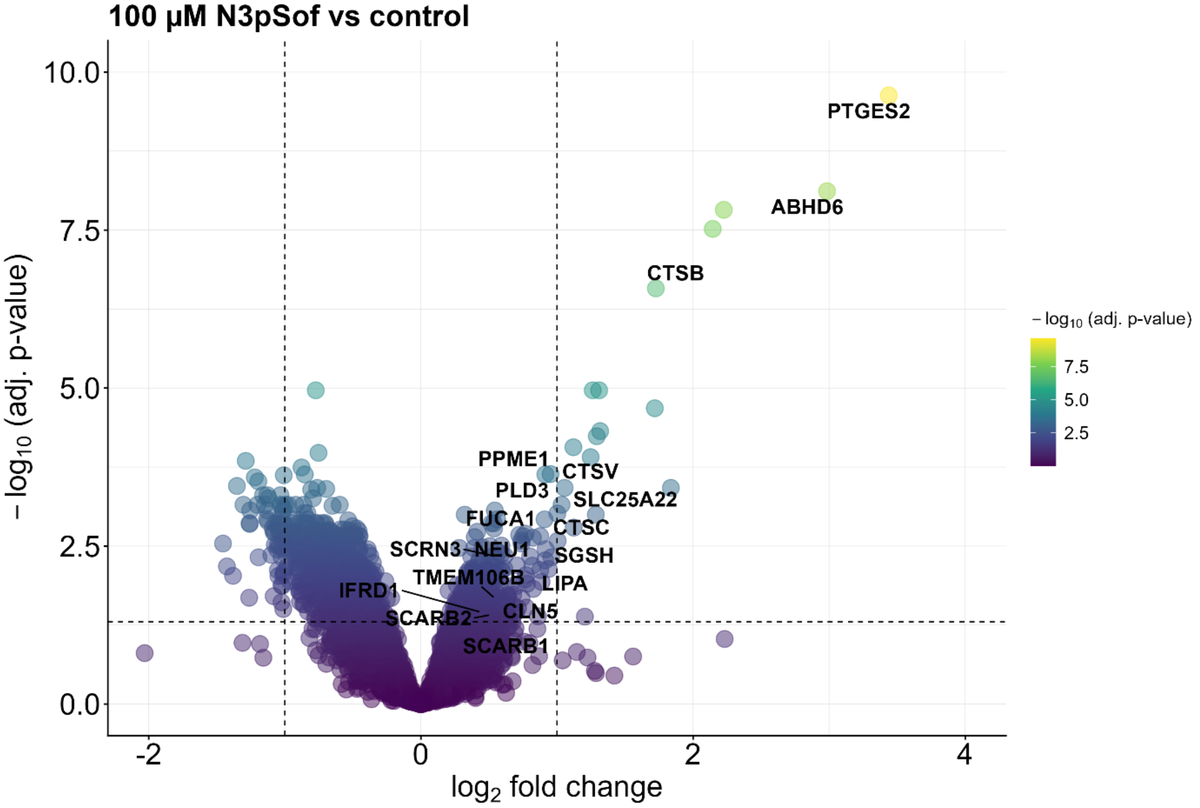

**Figure S2. Protein enrichment by N3pSof in HEK293T cells.** Volcano plot visualizing the proteins enriched by 100 µM N3pSof probe (*n* = 4 biological replicates, cut-off lines: adj. *p*-value = 1.3, fold change = 1). Labelled are all proteins that are enriched in either the 100 µM or 200 µM N3pSof treated group or the proN6pA treated group (overlapping proteins in main figure 2 C)).

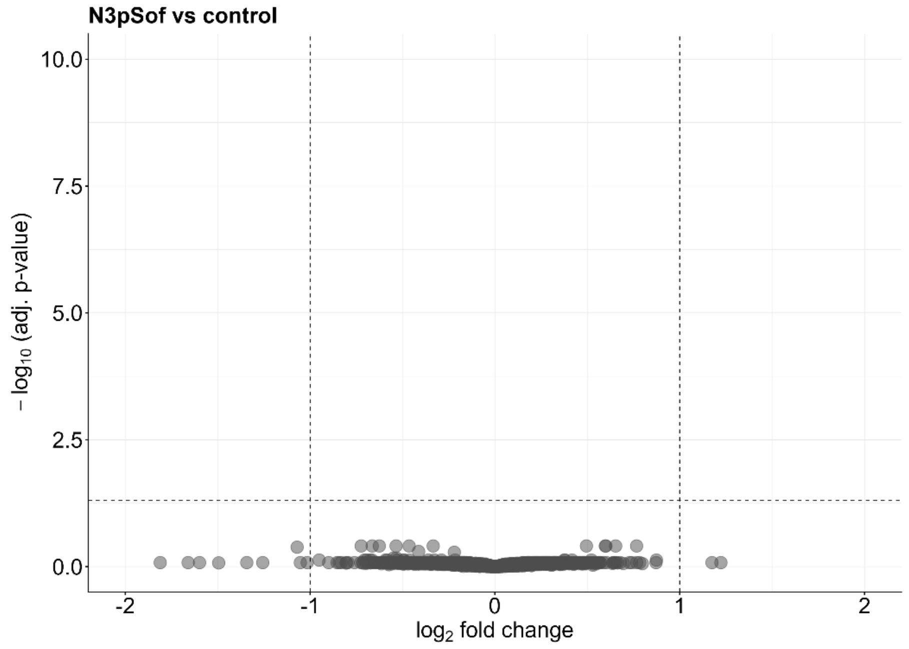

**Figure S3. Whole proteome analysis of N3pSof treated HEK293T cells**. Cells were treated with 100 µM N3pSof or DMSO as a control for 16 h. (*n* = 4 biological replicates, cut-off lines: adj. *p*-value = 1.3, fold change = 1).

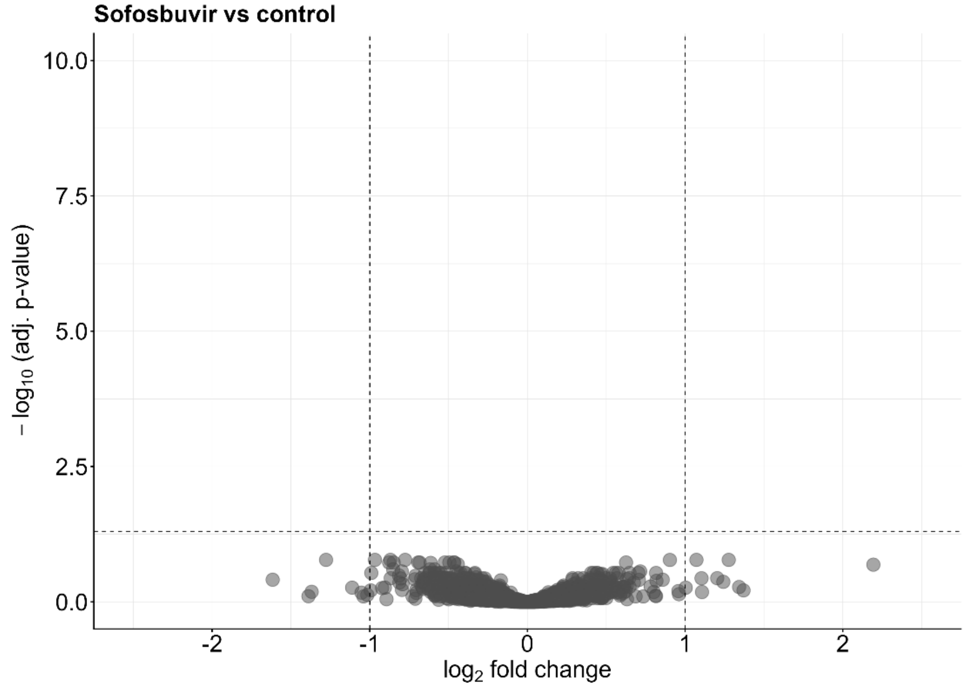

**Figure S4. Whole proteome analysis of Sofosbuvir treated HEK293T cells**. Cells were treated with 100 µM Sofosbuvir or DMSO as a control for 16 h. (*n* = 4 biological replicates, cut-off lines: adj. *p*-value = 1.3, fold change = 1).

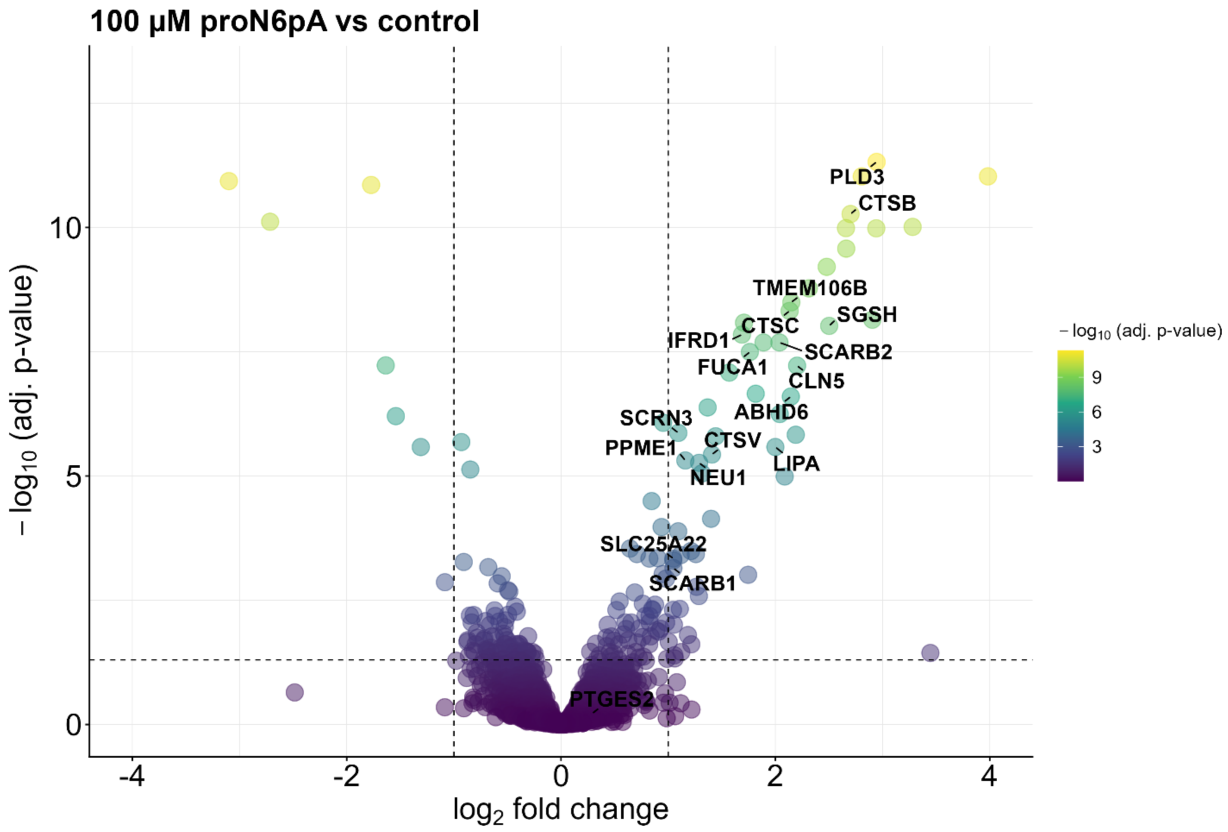

**Figure S5. Protein enrichment by proN6pA in HEK293T cells.** Volcano plot visualizing the proteins enriched by 100 µM proN6pA probe (*n* = 4 biological replicates, cut-off lines: adj. *p*-value = 1.3, fold change = 1). Labelled are all proteins that are enriched in either the 100 µM or 200 µM N3pSof treated group or the proN6pA treated group (overlapping proteins in main figure 2 C)).

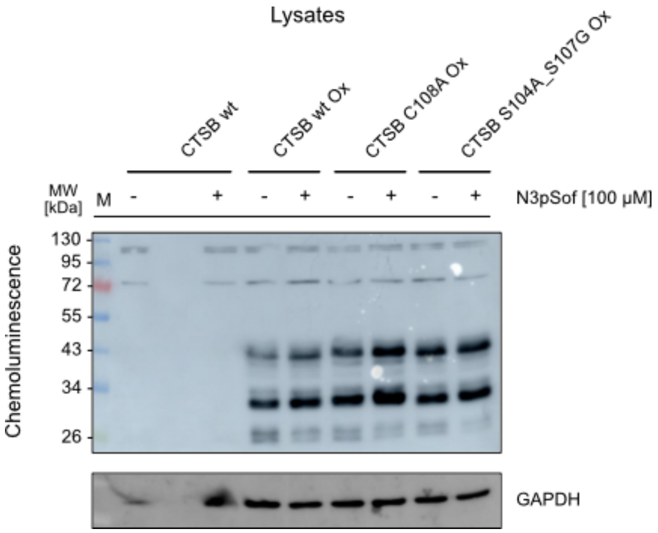

**Figure S6. Western blot of HEK293T whole cell lysates.** Western Blot showing lysates from wild-type cells or cells transiently overexpressing either wild-type CTSB, CTSB_C108A, or CTSB_S104A_S107G.

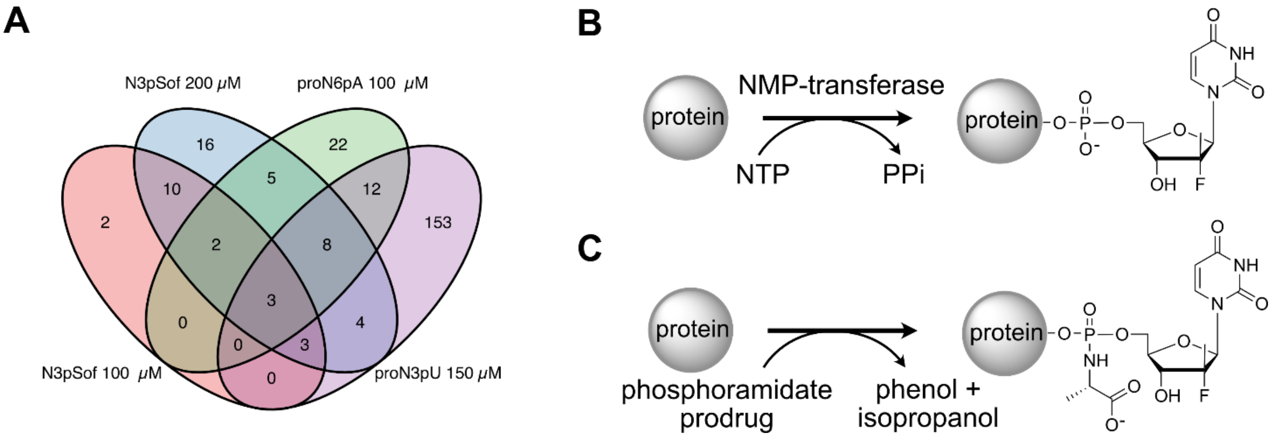

**Figure S7. Overlap and possible routes of protein enrichment by N3pSof, proN6pA, and N3pU. (A)** Overlap of enriched proteins between N3pSof and proN6pA (this study) and N3pU (Ref. [1]). **(B)** Probe incorporation by NMPylating enzymes. **(C)** Probe incorporation through chemical labelling.

#### Supplementary Tables

**Table S1.** Conditions for PCR cycle. The annealing temperature $T_a$ for each primer pair was determined based on its sequence.

| Step | Temperature | Time |
|---|---|---|
| Initial Denaturation | 98 °C | 30 seconds |
| 25 Cycles | 98 °C | 10 seconds |
| | $T_a$ | 30 seconds |
| | 72 °C | 30 seconds/kb |
| Final Extension | 72 °C | 2 minutes |
| Hold | 4 °C | |

**Table S2.** Sequences of oligonucleotides (5' to 3') used for sequencing.

| Construct | Sense | Sequence (5' to 3') |
|---|---|---|
| CTSB C108A | Forward | ACACCAATGCGCACGTCA |
| CTSB S104A S107G | Forward | TAATACGACTCACTATAGGG |

**Table S3.** Sequences of oligonucleotides (5' to 3') used for cloning. $T_a$ - annealing temperature.

| Protein | Mutant | Sense | Sequence (5' to 3') | $T_a$ [°C] |
|---|---|---|---|---|
| CTSB | C108A | Fwd | CTGTGGCAGCGCGTGGGCCTTCG | 83.1 |
| | | Rev | GAGCCCTGGTCTCTG | 54.4 |
| CTSB | CTSB S104A S107G | Fwd | TCAGAGACCAGGGCGCCTGTGGCGGCTGCTGGGCC | 72 |
| | | Rev | TCTCTTTGATGGTGGGACACTGTGGCCATTGTTCCCGT | 72 |

#### Organic Synthesis

Reagents and solvents were purchased from commercial suppliers, including abcr GmbH (Karlsruhe, Germany), Acros Organics (Thermo Fisher Scientific, USA), Sigma-Aldrich (St. Louis, USA), TCI Deutschland GmbH (Eschborn, Germany), Thermo Scientific (Loughborough, UK) and they were used without any further purification.

TLC (thin layer chromatography) was performed to monitor the reaction progress, which was done on precoated silica gel plates (60 F-254, 0.25 mm, from Merck KGaA, Darmstadt, Germany) as the stationary phase, with detection by potassium permanganate staining and gentle heating.

Flash chromatography was performed on Pure Chromatography Instruments (Pure-C815 Flash) from BUCHI Labortechnik GmbH (Essen, Germany) on pre-fabricated FlashPure EcoFlex Silica columnsmwith the indicated eluents.

$^{1}$H and proton-decoupled $^{13}$C NMR spectra for compound characterization in deuterated chloroform ($CDCl_3$) were acquired at 298K on a Bruker Avance Neo 500 spectrometer (11.7 T), using a nitrogencooled Prodigy BBO probe. All chemical shifts are reported in delta (δ) units in parts per million (ppm) relative to distinguished solvent signals as an internal reference. Coupling constant *J* are indicated in Hertz (Hz). Splitting patterns for peak assignments are indicated as the following abbreviations: s, singlet; d, doublet; t, triplet; q, quartet; m, multiplet; dd, doublet of doublets; dt, doublet of triplets. Spectra were analyzed using MestReNova (version 14.2.1, Mestrelab Research, Santiago de Compostela, Spain).

LC-MS chromatograms were obtained from Thermo Fisher LC-MS system composed of a DIONEX UltiMate 3000 HPLC system (pump, auto sampler, column compartment, and diode array detector) and an ESI (electrospray ionization)-MS based MSQ Plus single quadrupole mass spectrometer; the latter one was used for ESI-MS measurements with direct injection. Reversed phase column chromatographic methods use a Hypersil Gold C18 selectivity column (100 × 2.1 mm).

**N3-propargyl-Sofosbuvir**

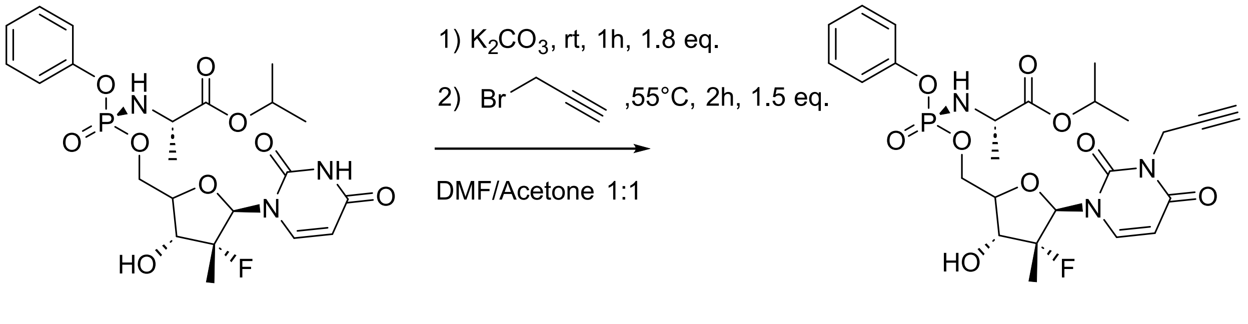

**Sofosbuvir** **N3-propargyl-Sofosbuvir (N3pSof)**

67%

To a solution of Sofosbuvir (1g, 1.89 mmol, 1 eq.) in a mixture of Acetone and DMF (1:1) was added finely ground $K_2CO_3$ (469.9 mg, 3.4 mmol, 1.8 eq.). The suspension was stirred at room temperature for 1h after short sonification to resolve all clumps. Propargyl bromide was added dropwise. The reaction mixture was stirred at 55°C for 2h. Subsequently the suspension was filtered. The solvents were removed under reduced pressure and then co-evaporated with toluene (3x 5 mL). The crude product was purified by column chromatography on silica (DCM/MeOH 5%). N3-propargyl-Sofosbuvir (715 mg, 1.26 mmol, 67%) was obtained as colorless oil.

$^{1}$H NMR (500 MHz, $CDCl_3$) δ 7.45 (d, *J* = 8.2 Hz, 1H), 7.38 – 7.30 (m, 2H), 5.77 (d, *J* = 8.2 Hz, 1H), 5.07 – 4.96 (m, *J* = 6.1 Hz, 1H), 4.69 (d, *J* = 2.5 Hz, 2H), 4.53 (ddd, *J* = 12.5, 8.1, 2.0 Hz, 1H), 4.44 (ddd, *J* = 12.4, 8.5, 2.5 Hz, 1H), 4.11 (dt, *J* = 9.4, 2.0 Hz, 1H), 3.99 – 3.80 (m, 2H), 2.95 (s, 1H), 2.88 (s, 1H), 2.17 (t, *J* = 2.5 Hz, 1H), 1.75 (s, 4H), 1.46 – 1.30 (m, 6H), 1.24 (dd, *J* = 6.3, 1.3 Hz, 7H).

$^{13}$C NMR (126 MHz, $CDCl_3$) δ 172.91, 172.86, 162.47, 161.05, 150.32, 150.27, 150.11, 129.86, 129.82, 125.25, 119.68, 119.64, 102.30, 100.95, 99.50, 79.72, 77.61, 77.16, 77.11, 76.91, 76.65, 71.78, 71.64, 70.67, 69.63, 69.56, 63.85, 50.29, 36.37, 31.34, 30.15, 21.62, 21.50, 20.77, 20.73, 16.46, 16.26, 0.90.

HR-MS (ESI+): m/z $[M+H]^+$ calculated for $C_{25}H_{32}FN_3O_9P$: 568,1855, found: 568,1846

HR-MS (ESI-): m/z $[M-H]^-$ calculated for $C_{25}H_{30}FN_3O_9P$: 566,1709, found: 566,1701

#### Analytical and Biochemistry methods

##### Culturing of HEK293T cells

HEK293T (CVCL_0063) cells were cultured in Dulbeccos Modified Eagles Medium – high glucose (DMEM) supplemented with 10% fetal bovine serum (FBS) and 2 mM L-alanyl-L-glutamine at 37 °C and 5% $CO_2$ atmosphere.

##### Probe treatment *in cellulo*

Cells were either seeded with 1 * $10^6$ cells in p60, 2 * $10^6$ cells in p100 or 4 * $10^6$ cells in p150 dishes in 5 mL, 10 mL or 20 mL of culture medium. Cells were either treated with N3pSof (100 – 1000 µM, varying by experiment) or respective amount of DMSO as control. After probe addition, cells were incubated 16 h prior to harvest. Cells were harvested by washing twice with 4 or 8 mL cold DPBS (4 °C), scraped in 0.5 mL (p60) or 1 mL (p100) or 2 mL (p150) DPBS and centrifuged at 200 x g, 4 °C, 5 min. After removal of the supernatant, the cell pellets were frozen at -80 °C or directly subjected to cell lysis.

##### Cell lysis

Cells were lysed with 100 – 400 µL of lysis buffer (1% NP-40, 0.2% SDS, 20 mM Hepes, pH 7.5) via rod sonication (10 s, 20% intensity). The lysates were clarified by centrifugation (13,000 x g, 4 °C, 10 min) and supernatant was transferred into a new Eppendorf tube. The lysates were stored at -80 °C or processed directly.

##### Protein concentration measurement

Protein concentration measurement was performed with a Pierce™ BCA Protein Assay Kit (Thermo Scientific).

##### In-gel fluorescence analysis

100 µg protein per sample/replicate was diluted with lysis buffer (1% NP-40, 0.2% SDS, 20 mM Hepes, pH 7.5) ad 93.4 µL. 1 µL TAMRA-$N_3$ (10 mM in DMSO), 3 µL TCEP (100 mM in MS water) and 0.6 µL TBTA (16.7 mM in DMSO) were added and the sample was vortexed and shortly spun down. 2 µL 50 mM $CuSO_4$ was added to 100 µL final volume and the sample was vortexed, shortly spun down and incubated 1.5 h at rt, 650 rpm protected from light. Proteins were precipitated by the addition of 400 µL cold Acetone (-20 °C) and incubation at -20 °C for 1 h to O/N. The precipitated proteins were centrifuged for 10 min, 13.000 x g, 4 °C and the supernatant was removed. Proteins were reconstituted in 80 µL 0.2% SDS in PBS and sonicated 5 sec at 20% intensity with a rod sonicator. For SDS PAGE 20 µg protein per sample (16 µL) were mixed with 4 µL 5x Laemmli buffer + DTT (200 µL 5x LB + 20 µL 1M DTT) and heated for 5 min at 95 °C. Samples were loaded on a 10% self-made SDS-PAGE gel after having cooled down to rt, and the gel was run for 10 min at 100V and then 50 – 60 min at 150V in 1x running buffer (25 mM Tris, 0.192 M glycine, 0.1% (m/v) SDS). In-gel fluorescence was scanned with Amersham Imager 680 (GE Healthcare). For loading control, proteins were stained with a coomassie staining solution (0.25% Coomassie Blue R-250, 10% acetic acid, 50% MeOH) and properly destained with destaining solution (20% MeOH, 10% acetic acid).

##### Western blot analysis

After running the SDS PAGE the gel was washed 3x with deionized water and the stacking gel was removed. The membrane was equilibrated in MeOH for 1 min and then the gel, blotting papers and the membrane were equilibrated in semi dry transfer buffer (5.82 g Tris, 2.9 g Glycine, 0.375 g SDS, 20% MeOH in 1 L deionized water) for 10 min. The blotting sandwich was assembled, and proteins were blotted over 30 min at 25V. After complete blotting the membrane was incubated in blocking buffer (5% milk powder in PBS + 0.5% Tween 20) for 1 h at rt, shaking. Subsequently, the membrane was incubated O/N with 10 mL of the respective primary antibody (1:1000 in blocking buffer) at 4 °C, shaking. The next day, the membrane was washed 3x with PBS + 0.5% Tween 20 for 5 min and then incubated with HRP-coupled secondary antibody (1:10.000 in blocking buffer) and anti-GAPDH rhodamine coupled antibody (1:10.000 in blocking buffer) simultaneously for 1 h at rt, protected from light. The secondary antibody and anti-GAPDH antibody were removed, and the membrane was washed 3x 10 min with PBS + 0.5% Tween 20. Chemiluminescence was scanned with Amersham Imager 680 (GE Healthcare) after activation of the membrane with Amersham ECL Prime Western Blotting Detection Reagent (1:1).

##### Polymerase Chain Reaction (PCR)

PCR was performed for cloning purposes using plasmid DNA as a template. Here, the procedure was performed according to the manufacturers' protocol (Q5 site–directed mutagenesis, NEB

#E0554S). The thermocycling conditions applied are listed in Table S1. PCR product formation was validated by 0.7 % agarose gel electrophoresis.

##### 0.7% Agarose Gel

For validation of PCR product formation, 1% agarose gel electrophoresis was conducted. The agarose was dissolved in 40 ml 1x 40 mM Tris, 20 mM acetic acid, 1 mM EDTA buffer, and heated up in the microwave. After cooling down, 2 µl Gel stain was added and the gel was polymerized. As a standard, 1 kB DNA ladder (Carl Roth) was used. To the PCR product, 6x stain (NEB) was added (cf = 1x). The electrophoresis was conducted for 1 h at 100 V in 0.5x Tris, acetic acid, and EDTA buffer. PCR products were detected under UV light.

##### KLD Reaction

A KLD (Kinase, Ligase, DpnI) reaction was performed with the PCR product according to the manufacturers' protocol (Q5 site–directed mutagenesis, NEB #E0554S). Here, the PCR product is first phosphorylated (kinase), then ligated (ligase), and residual template DNA is decomposed by DpnI enzyme (only methylated DNA is a substrate). The reaction mixture was prepared, mixed by pipetting up and down 5 to 10 times, and incubated for 5 min at room temperature. The KLD mixture was used for subsequent bacterial transformation.

##### Chemical Transformation of Bacterial Cells

For chemical transformation, chemically competent E. coli DH5-alpha cells stored at −80 °C were thawn for 10 min on ice. Then, 5 µl of the Kinase, Ligase, and DpnI (KLD) reaction mixture was added directly to 50 µl of cells, and the tube was gently flicked three times. The cells were incubated for 30 min on ice and then heat shocked at 42 °C for 30 s. The cells were immediately placed on ice for 5 min for regeneration before the addition of 950 µl of SOC medium (NEB) and incubation for 1 h at 37 °C at 250 rpm. In the meantime, LB-Agar (1.3% Agar) plates supplemented with 100 µg/ml ampicillin were prewarmed at 37 °C. After incubation, 100 µl of the cell suspension was spread onto prewarmed LB-Agar plates and evenly distributed using glass beads by shaking the plate under sterile conditions. The LB-Agar plates were then incubated overnight at 37 °C and subsequently stored at 4 °C or directly used for plasmid amplification.

##### Q5 Site–Directed Mutagenesis

For preparation of hCTSB C108A and S104A_S107G mutants starting from pcDNA3.1_hCTSB (RRID:Addgene_11249), the Q5 Site-Directed Mutagenesis Kit (New England Biolabs #E0554S) was used. The sequence was validated by Sanger sequencing (Genewiz) using the sequencing oligonucleotides listed in Table S2. Sequences of primers used for the introduction of the mutations are shown in Table S3.

**Plasmid DNA Isolation Using Mini/Midi Preparation Kit**

For plasmid DNA isolation, either the Mini (small scale, NEB) or Midi (large scale, Zymo) preparation kit was utilized. For Mini or Midi preparation, 3 ml or 100 ml, respectively, of sterile LB medium supplemented with 100 µg/ml ampicillin was prepared. A single clone from the overnight LB agar plate was selected using a pipette tip and subsequently added to the medium. The medium was then incubated for 12 to 16 h at 37 °C at 180 rpm. Plasmid DNA was isolated following the manufacturer's protocol for the Plasmid Miniprep/Midiprep Kit. The DNA was eluted in the respective elution buffer included in the kit. DNA concentration was measured using a Nanodrop, and the DNA sequence was confirmed by Sanger sequencing conducted by Genewiz.

**Enrichment of modified proteins using the SP2E protocol for LC-MS/MS**

400 µg protein per sample/replicate was diluted with lysis buffer (1% NP-40, 0.2% SDS, 20 mM Hepes, pH 7.5) add 186.8 µL. 2 µL Biotin-$N_3$ (10 mM in DMSO), 6 µL TCEP (100 mM in MS water) and 1.2 µL TBTA (16.7 mM in DMSO) were added and the sample was vortexed and shortly spun down. 4 µL 50 mM $CuSO_4$ was added to 200 µL final volume and the sample was vortexed, shortly spun down and incubated for 1.5 h at r.t., 650 rpm. After the incubation was completed, 200 µL of 8 M urea in MS-water was added to the click reaction to a final volume of 400 µL. 100 µL of a 1:1 mixture of hydrophobic and hydrophilic carboxylate-coated magnetic beads (Cytiva) was washed thrice with 500 µL of MS-water and the reaction mixture was transferred onto the pre-washed beads followed by the addition of 600 µL absolute Ethanol, mixing and incubation at r.t., 950 rpm for 5 min. The beads were washed thrice with 500 µL of 80% EtOH in MS-water. Subsequently, proteins were eluted from the carboxylate-coated beads twice with 500 µL of 0.2% SDS in PBS at r.t., 950 rpm for 5 min and transferred onto 50 µL streptavidin-coated magnetic beads (New England Biolabs) that had been equilibrated 3x with 500 µL 0.2% SDS in PBS. For streptavidin-biotin complex formation, beads were incubated at r.t., 950 rpm for 1 h with eluted proteins. After the incubation, beads were washed three times with 500 µL of 0.1% NP-40 in PBS, two times with 500 µL of 6 M urea (in MS-water) and two times with 500 µL of MS-water by vortexing and short-spin between each wash step. After the last wash step, 80 µL of ABC buffer (ammoniumbicarbonate, 125 mM in MS-water), 10 µL of TCEP (100 mM in MS-water) and 10 µL of chloroacetamide (400 mM in MS-water) was added and incubated at 95 °C for 5 min. Then, proteins were digested O/N with 1.5 µL sequencing-grade trypsin (0.5 mg/mL, Promega) at 37 °C, 650 rpm. The next day supernatants were transferred into new Eppendorf tubes, and beads were washed thrice with 100 µL ABC buffer (100 mM in MS-water) to collect the peptides (wash fractions were combined into the corresponding tubes). The combined fractions were acidified with 2 µL of MS-grade FA and peptides were desalted with 50 mg of SepPak C18 cartridges on a vacuum manifold. First, cartridges were equilibrated with 1 mL of MS grade ACN, 1 mL of elution buffer (80% ACN with 0.5% FA in MS-water) and 3 mL of wash buffer (0.5% FA in MS-water). Second, combined peptide fractions were loaded onto the cartridges and washed with 3 mL of wash buffer. Finally, peptides were eluted twice with 250 µL of elution buffer. Desalted peptide eluates were vacuum-dried with a SpeedVac at 35 °C, reconstituted in 30 µL of 1% FA in MS-water (vortex and 15 min sonication bath), transferred to MS vials and subjected to LC-MS/MS analysis with 2 – 5 µl injection volume.

#### Enrichment of modified proteins using the SP2E protocol for SDS PAGE & Western Blot

300 µg protein per sample/replicate was diluted with lysis buffer (1% NP-40, 0.2% SDS, 20 mM Hepes, pH 7.5) add 93.4 µL. 1 µL TAMRA-Biotin-$N_3$ (10 mM in DMSO), 3 µL TCEP (100 mM in MS water) and 0.6 µL TBTA (16.7 mM in DMSO) were added and the sample was vortexed and shortly spun down. 2 µL 50 mM $CuSO_4$ was added to 100 µL final volume and the sample was vortexed, shortly spun down and incubated for 1.5 h at r.t., 650 rpm, protected from light. After the incubation was completed, 100 µL of 8 M urea in MS-water was added to the click reaction to a final volume of 200 µL. 100 µL of a 1:1 mixture of hydrophobic and hydrophilic carboxylate-coated magnetic beads (Cytiva) was washed thrice with 500 µL of MS-water and the reaction mixture was transferred onto the pre-washed beads followed by the addition of 600 µL absolute Ethanol, mixing and incubation at r.t., 950 rpm for 5 min. The beads were washed thrice with 500 µL of 80% EtOH in MS-water. Subsequently, proteins were eluted from the carboxylate-coated beads twice with 500 µL of 0.2% SDS in PBS at r.t., 950 rpm for 5 min and transferred onto 50 µL streptavidin-coated magnetic beads (New England Biolabs) that had been equilibrated 3x with 500 µL 0.2% SDS in PBS. For streptavidin-biotin complex formation, beads were incubated at r.t., 950 rpm for 1 h with eluted proteins. After the incubation, beads were washed three times with 500 µL of 0.1% NP-40 in PBS, two times with 500 µL of 6 M urea in MS-water and two times with 500 µL of MS-water by vortexing and short-spin between each wash step. After the last wash step, the beads were resuspended in 50 µL 1x Laemmli Buffer supplemented with 20 mM DTT and incubated for 5 min at 95 °C. The beads were removed on a magnetic rack, and the supernatant was transferred into a new Eppendorf tube. For unprocessed lysates 20 µg protein was diluted in 0.2% SDS in PBS ad 16 µl and 4 µL 5x Laemmli Buffer + 100 mM DTT was added (final 1x LB + 20 mM DTT) and incubated for 5 min at 95 °C. 20 µL of enriched or unprocessed sample was loaded onto a 10 % SDS gel. The PAGE was run at 100V for 10 min, and then at 150V for ~60 min, while protected from light. Subsequently, Western blot analysis was performed as previously described.

#### Enrichment of modified peptides using desthiobiotin-tag (Site-ID)

Probe or DMSO treated cells were harvested by washing twice with 8 mL cold DPBS (4 °C), scraped in 3 mL cold DPBS and then 3x 1 mL of this suspension was split in Eppendorf tubes, yielding 3 technical replicates of each probe treated and DMSO treated samples. Each replicate was lysed in 1300 µL lysis buffer (1% NP-40, 0.2% SDS, 20 mM Hepes, pH 7.5) as previously described and protein concentration was determined by Pierce™ BCA Protein Assay Kit (Thermo Scientific). A total amount of 8 - 10 mg protein of each control and *in cellulo* probe-treated lysates (split across three replicates each) was used for the subsequent workflow. All replicates were diluted with lysis buffer ad 1893.5 µL. 4 µL Desthiobiotin-$N_3$ (50 mM in DMSO), 60 µL TCEP (100 mM in MS water) and 2.5 µL TBTA (83.5 mM in DMSO) were added, and the sample was vortexed and shortly spun down. 40 µL 50 mM $CuSO_4$ was added to 2 mL final volume, and the sample was vortexed, shortly spun down and incubated 2.5 h at rt, 650 rpm. Subsequently, the click reaction mixture was transferred into a 15 ml falcon tube, 8 mL cold acetone (-20 °C) was added and proteins were precipitated at -20 °C O/N. Precipitated proteins were spun down 10 min, 13,000 x g, 4 °C, supernatant was disposed, and each protein pellet was washed twice with 1 mL of cold methanol by resuspending the pellet with sonication (3x, 5 s, 20% intensity) and centrifugation. After the last wash step, methanol was removed, and the pellet was air-dried for 10 min. The pellet was reconstituted in 300 µL of 8 M urea in 0.1 M TEAB solution by sonication (3x, 10 s, 20% intensity) and diluted with 900 µL of 0.1 M TEAB ad 1.2 mL total volume. 100 µL of streptavidin-coated agarose beads (Sigma Aldrich) per sample (probe and control) were pre-washed thrice with 1 mL of 0.2% NP-40 in PBS. The washed beads were split equally across 2x three Eppendorf tubes (3x probe, 3x control). The reconstituted replicates were added to the beads and incubated for 1.5 h at rt, 850 rpm. All the subsequent washing steps were performed by shortly vortexing the tubes, centrifuging 2 min, 2000 rpm, r. t. and disposal of the supernatant. For enrichment, the beads were washed thrice with 1 mL 0.1% NP-40 in PBS (the three replicates of each sample were combined after the first washing step), thrice with 1 mL of PBS, thrice with 1 mL of MS-grade $H_2O$ and once with 1 mL 2 M urea in 100 mM ABC buffer. After the last supernatant was discarded, the beads were resuspended in 190 µL 2 M urea in 100 mM ABC buffer, 2 µL of 1 M DTT was added and incubated at 37 °C for 45 min, 1000 rpm. The reduced thiol groups were alkylated by the addition of 8 µL 0.5 M iodoacetamide (fresh prepared with 2 M urea in 100 mM ABC buffer) and incubation at r. t. for 1 h. 600 µL 100 mM ABC buffer was added, vortexed, centrifuged at 1000 x g, 2 min and the supernatant was removed. For digestion, beads were resuspended in 100 µL 100 mM ABC buffer, then 4 µL of sequencing-grade trypsin (0.5 mg/mL, Promega) was added and proteins were digested at 37 °C, 1000 rpm, O/N. The next day, 800 µL 0.1% NP-40 in PBS was added to the tryptic digest, centrifuged and supernatant was removed. The beads were washed thrice with 1 mL of 0.1% of NP-40 in PBS, thrice with 1 mL PBS and thrice with 1 mL MS grade $H_2O$. Desthiobiotin-modified peptides were eluted twice with 200 µL of ACN/$H_2O$ (1:1) with 0.1% FA, twice with 100 µL of ACN/$H_2O$ (1:1) with 0.5% FA and twice with 100 µL of ACN/$H_2O$ (4:1) with 0.5% FA. For each step, beads were incubated with the corresponding elution buffer for 10 min, 1000 rpm, 40 °C, spun down and the supernatant eluate fraction collected and combined. Combined fractions were centrifuged for 1 min, 20,000 x g and transferred into a new Eppendorf tube while ~50 µL of the eluate was left to ensure that no beads were collected. The eluate was vacuum-dried with a SpeedVac at 35 °C, reconstituted in 20 µL of 1% FA in $H_2O$, vortexed and sonicated for 15 min. Finally, peptide mixtures were transferred to MS-vials and submitted to LC-MS/MS measurement with an injection volume of 8 µL.

**Whole proteome analysis using the SP3 protocol**

10 µg protein per replicate was diluted with lysis buffer (1% NP-40, 0.2% SDS, 20 mM Hepes, pH 7.5) add 10 µL in a 96 well plate. A 1:1 mixture of hydrophobic and hydrophilic carboxylate-coated magnetic beads (Cytiva) was washed thrice with 500 µL of MS-water and resuspended to the initial volume. 2 µl of these pre-washed beads were added to the proteins, followed by the addition of 48 µL absolute Ethanol and 10 min incubation at rt, 1000 rpm. The following washing steps were performed by the Microlab prep pipetting robot (Hamilton). The 96 well plate was transferred to a magnetic rack and incubated for 2 min. The supernatant was discarded. Beads were washed twice with 180 µL 80% EtOH (30s incubation, rt, shaking). Beads were washed once with 180 µL of ACN (LC-MS grade, 30s incubation, rt, shaking). Supernatant was discarded and beads were air-dried for 30s. The beads were reconstituted in 20 µL 100 mM ABC buffer and 1 µL trypsin (0.5 mg/mL, Promega) was added and proteins were digested at 37 °C, 1000 rpm, O/N. The next day beads were resuspended by pipetting, and the supernatant was removed on a magnetic rack and transferred into new Eppendorf tubes. Proteins were eluted twice by the addition of 10 µL MS-water and 5 min incubation at 40 °C. 4 µL 1% formic acid in MS water was added to the combined fractions, tubes were placed on a magnetic rack and peptides were transferred into MS vials while ~ 20 µL were left to ensure that no beads were collected. Samples were subjected to LC-MS/MS analysis with 1 µL injection volume.

#### LC-MS/MS measurement

MS measurements were performed on a Orbitrap Eclipse Tribrid Mass Spectrometer (Thermo Fisher Scientific) coupled to an UltiMate 3000 Nano-HPLC (Thermo Fisher Scientific) via a Nanospray Flex (Thermo Fisher Scientific) and FAIMS interface (Thermo Fisher Scientific). First, peptides were loaded on an Acclaim PepMap 100 µ-precolumn cartridge (5 µm, 100 Å; 300 µM ID x 5 mm, Thermo Fisher Scientific). Then, peptides were separated at 40 °C on a PicoTip emitter (noncoated, 15 cm, 75 µm ID, 8 µm tip, New Objective) that was in house packed with ReprosilPur 120 C18-AQ material (1.9 µm, 150 Å, Dr. A. Maisch GmbH) or on an Aurora Elite XT C18 UHPLC column (15 cm, 75 µm ID, IonOpticks). The LC buffers consisted of MS-grade water (A) and acetonitrile (B) both supplemented with 0.1% formic acid. The gradient was run from 4-35.2% B during a 60 min method (0-5 min 4%, 5-6 min to 7%, 7-36 min to 24.8%, 36-41 min to 35.2%, 40-41.1 min 80%, 41.1-46 min 80%, 46-60 min 4%) at a flow rate of 300 nL/min.

*Data-independent acquisition for SP2E enrichment*

FAIMS was performed with one CV at -45 V. One DIA cycle comprised one MS1 scan followed by 30 MS2 scans. The mass spectrometer was operated in DIA mode with following settings: Polarity: positive; MS1 Orbitrap resolution: 60k; MS1 AGC target: standard; MS1 maximum injection time: 50 ms; MS1 scan range: m/z 200-1800; RF Lens: 30%; Precursor Mass Range: m/z 500-740; isolation window: m/z 4; window overlap: m/z 2; MS2 Orbitrap resolution: 30k; MS2 AGC target: 200%; MS2 maximum injection time: auto; HCD collision energy: 35%; RF Lens: 30%; MS2 scan range: auto.

*Data-dependent acquisition for Desthiobiotin enrichment (Site-ID)*

The mass spectrometer was operated with FAIMS in DDA mode with following settings for MS1: FAIMS CV: -50 V and -70 V; Polarity: positive; Orbitrap Resolution: 240k; Mass Range: Normal; Scan Range: 375 – 1500; RF Lens: 30%; AGC Target: Standard; Maximum Injection Time Mode: Custom (35 ms); Microscans: 1; Intensity Threshold: 1.0e4; Included charge state(s): 2-6; Dynamic Exclusion: 60s; Mass Tolerance: 10 ppm. Cycle Time: 1.7s; MS2 spectra were acquired with following settings: Orbitrap Resolution: 15k; Mass Range: Normal; Scan Range Mode: Auto; AGC Target:100%; Maximum Injection Time Mode: Custom (35 ms); HCD Collision Energy: 30%.

#### Computational evaluation of measured mass spectra

DIA-NN (peptide quantification)

Measured .raw files were analysed with DIA-NN 2.1 and peptides were searched against Uniprot database for Homo sapiens (UP000005640, taxon identifier: 9606) with included isoforms The DIA-NN[2] settings were as follows: Contaminants: enabled; FASTA digest for library-free search/library generation: enabled; Deep learning-based spectra, RTs and IMs prediction: enabled; missed cleavages: 2; max number of variable modifications: 2; modifications: N-term M excision, carbamidomethylation, oxidation (M) and N-term acetylation; Peptide length range: 7 – 35; Precursor charge range: 2-6; precursor range: m/z 500-740; fragment ion range: m/z 200-1800; precursor FDR level: 1%; match between runs (MBR): enabled; all other parameters were used as default.

FragPipe platform (Site-ID)

MS *.raw files were converted to *.mzML format with “MSConvert” with following settings: “peakPicking” filter with “vendor msLevel = 1”, “titleMaker” filter was enabled. Peptides were searched in Fragpipe 23.1 against Uniprot database for Homo sapiens (UP000005640, taxon identifier: 9606) with included contaminants and decoys. Due to the huge variety of settings, the four general FragPipe workflow (open search) is provided on the PRIDE server. In the following most important features were listed:

*Closed search:* variable modifications: 774.31134(STYC), carbamidomethylation (C), 845.34845 (STYC); fixed modifications: none

##### Data statistics and evaluation from computational output

Statistics and plots were generated using a custom R script. In brief, the “report.parquet” file obtained from the DIA-NN calculation was loaded as the main data using the readDIANN function from the limpa package[3] with q.cutoffs set to 0.01. Next, non-proteotypic peptides were filtered out and groups were added to specify control or probe treated samples. The detection-probability curve was generated using the dpc function and proteins were quantified from the “Protein.Names” column using dpcQuant (for heatmaps proteins were quantified from the “Genes” column to avoid multiple entries for a single gene)[3]. Differential expression analysis was performed using the limma package[4]. Volcano plots were generated using the ggplot2 package and cutoffs were set to 0.05 for the adjusted p-value (corrected using the Benjamin Hochberg method) and 0.01 for log2 fold change.

*Closed search for modified peptides (FragPipe)*: Mass shifts of modified peptides were obtained from the “psm.tsv” file “Assigned Modifications” column.

#### Materials

| Reagent/Resource | Reference or Source | Identifier or Catalog Number |
|---|---|---|
| **Experimental Models** | | |
| Human: HEK293T | Ref.[5] | RRID: CVCL_0063 |
| **Recombinant DNA** | | |
| pcDNA3.1_hCTSB | Hyeryun Choe Lab | RRID:Addgene_11249 |
| pcDNA3.1_hCTSB_C108A | This study | Derived from Addgene_11249 |
| pcDNA3.1_hCTSB_S104A_S107G | This study | Derived from Addgene_11249 |
| **Antibodies** | | |
| Anti-CTSB | proteintech | Cat# 12216-1-AP |
| Anti-GAPDH-Rhodamine | Bio-Rad | Cat# 12004168 |
| Goat-anti-rabbit HRP | Thermo Fisher | Cat# 31460 |

| Chemicals and Enzymes | | |
|---|---|---|
| Acetic acid (100%) | In house chemical store | N/A |
| Acetone (HPLC grade) | VWR Chemicals | Cat# 20067.320 |
| Acetonitrile (LC-MS grade) | Thermo Fisher | Cat# A955-212 |
| Agar-Agar | Carl Roth | Cat# 6494.3 |
| Alanyl-Glutamine | Sigma-Aldrich | Cat# G8541 |
| Ammonium Acetate | Sigma-Aldrich | Cat# 09689-1kg |
| Ammonium bicarbonate | Fluka | Cat# 09830-100g |
| Ammoniumperoxodisulfat | Sigma-Aldrich | Cat# 09913 |
| Ampicillin | Carl Roth | Cat# HP62.1 |
| Antibiotic-Antimycotic | Gibco | Cat# 15240062 |
| Avidin-agarose beads | Sigma-Aldrich | Cat# A9207-5ML |
| Benchmark$^{TM}$ fluorescence Protein Standard | Invitrogen | Cat# LC5928 |
| Biotin-PEG$_3$-N$_3$ | Carbosynth | Cat# FA34890 |
| BSA | AppliChem | Cat# A6588 |
| Carboxylate-coated magnetic beads (hydrophobic) | Cytiva | Cat# 65152105050250 |
| Carboxylate-coated magnetic beads (hydrophilic) | Cytiva | Cat# 45152105050250 |
| Chloroacetamide | Sigma-Aldrich | Cat# C0267-100g |
| Color Prestained Protein Standard (10-250 kDa) | New England Biolabs | Cat# P7719S |
| Coomassie Blue R-250 | Fluka | Cat# 27816 |
| $CuSO_4$ x 5 H2O | Acros | Cat# 10627162 |
| dd$H_2O$ (LC-MS grade) | Honeywell | Cat# 15665350 |
| Desthiobiotin-PEG$_3$-N$_3$ | Sigma-Aldrich | Cat# 902020-25mg |
| DMEM (1x) | Sigma-Aldrich | Cat# D6546 |
| DMF anhydrous | Sigma-Aldrich | Cat# 227056 |
| DMSO | Sigma-Aldrich | Cat# D4540 |
| DPBS (1x) | Sigma-Aldrich | Cat# D8357 |
| DTT | AppliChem | Cat# A2948,0025 |
| EDTA | BioChemica | Cat# A1103 |
| Ethanol (EtOH) | Merck | Cat# 34852 |
| Fetal bovine serum | Thermo Fisher | Cat# A3840001 |
| Fetal bovine serum (FBS) | Biochrom | Cat# S0115 |
| Formic Acid (LC-MS grade) | Thermo Fisher | Cat# A117 |
| GelStain | Carl Roth | Cat# 3865.1 |
| Glycerol | Sigma-Aldrich | Cat# G5516 |
| HCl | In house chemical store | N/A |
| Hepes | Carl Roth | Cat# HN77.5 |
| HEPES solution | Sigma-Aldrich | Cat# H0887-100ML |

| | | |
|---|---|---|
| Iodoacetamide (IAA) | Sigma-Aldrich | Cat# I6125-10g |
| LB medium | Carl Roth | Cat# 66693 |
| Methanol (LC-MS grade) | Thermo Fisher | Cat# A456 |
| NaOH | Sigma-Aldrich | Cat# S5881 |
| NP-40 | Sigma-Aldrich | Cat# 74385 |
| Opti-MEM$^{TM}$ I Reduced Serum Medium, GlutaMAX™ Supplement | Thermo Fisher | Cat# 51985-026 |
| Penicillin/Streptomycin | Gibco | Cat# 15-140-122 |
| Polyethylenimine (PEI) | Polyscience | Cat# 23966 |
| Rotiphorese.Gel 30 (37,5:1) | Carl Roth | Cat# 3029.1 |
| sodium dodecyl sulfate | AppliChem | Cat# A2572 |
| Sofosbuvir | aablocks | Cat# AA000P0P |
| Streptavidin magnetic beads | New England Biolabs | Cat# S1420S |
| TAMRA-$N_3$ | Baseclick | Cat# BCFA-008-1 |
| TBTA | TCI | Cat# T2993 |
| TCEP | Carbosynth | Cat# FT01756 |
| TEAB (1 M) | Sigma-Aldrich | Cat# T7408 |
| TEMED | Sigma-Aldrich | Cat# T9281 |
| Tris-base | Thermo Fisher | Cat# 10724344 |
| Triton X-100 | Sigma-Aldrich | Cat# X100 |
| Trypan Blue | Thermo Fisher | Cat# 11538886 |
| TrypLE Express | Thermo Fisher | Cat# 12604013 |
| Trypsin (sequencing-grade) | Promega | Cat# V5113 |
| Tween 20 | Sigma-Aldrich | Cat# P6585 |
| Urea | AppliChem | Cat# A1049 |
| **Software** | | |
| FragPipe | Ref.[6–11] | https://fragpipe.nesvila b.org/ |
| DIA-NN | Ref.[2] | https://github.com/vde michev/DiaNN |
| MSConvert | Ref.[12] | https://proteowizard.so urceforge.io/download. html |
| PDV viewer | Ref.[13] | https://github.com/wen bostar/PDV |
| FreeStyle | Thermo Fisher Scientific | N/A |
| R | N/A | https://www.R-project.org/>. |
| RStudio | N/A | http://www.posit.co/ |
| **Kits and Other** | | |
| Aurora Elite XT C18 UHPLC column | ionopticks | Cat# AUR4-15075C18-XT |
| Pierce® BCA Protein Assay Kit | Thermo Fisher Scientific | Cat# 23225 |
| ZymoPURE™ II Plasmid Midiprep Kit | Zymo Research | Cat# D4200 |
| Plasmid Miniprep Kit | New England Biolabs | Cat# T1010S |

| | | |
|---|---|---|
| Q5 Site-Directed Mutagenesis Kit | New England Biolabs | Cat# E0554S |
| Orbitrap Eclipse Tribrid Mass Spectrometer | Thermo Fisher Scientific | N/A |
| FAIMS Pro Duo Interface | Thermo Fisher Scientific | N/A |
| PicoTip™ Emitter, Silica Tip™ | New Objectives | FS360-75-8-N-20-C15 |
| ReproSil-Pur 120 C18-AQ, 1.9 µm | Dr. Maisch GmbH | Cat# r119.aq.0001 |
| ReproSil-Pur 120 NH2, 3 µm | Dr. Maisch GmbH | Cat# r13.a0.0001 |
| PEPMAP100 C18 5UM 0.3X5MM | Thermo Fisher Scientific | Cat# 160454 |
| Sep-Pak C18 1 cc Vac Cartridge, 50 mg | Waters | 186000308 |
| VP 250/10 Nucleodur 100-5 C18 | AppliChem | Cat# 762022.100RC |

### NMR Spectra

*N*[3]-propargyl-Sofosbuvir

$^{1}H$ NMR (500 MHz, $CDCl_3$)

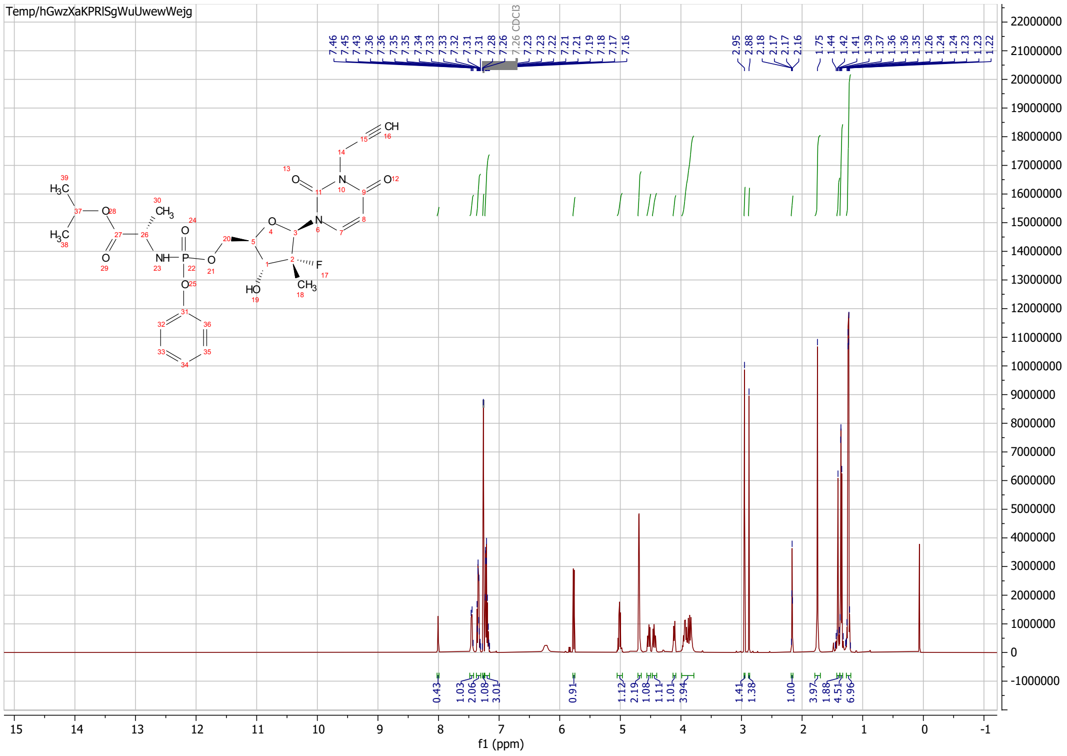

$^{13}C$ NMR {$^{1}H$} (126 MHz, $CDCl_3$)

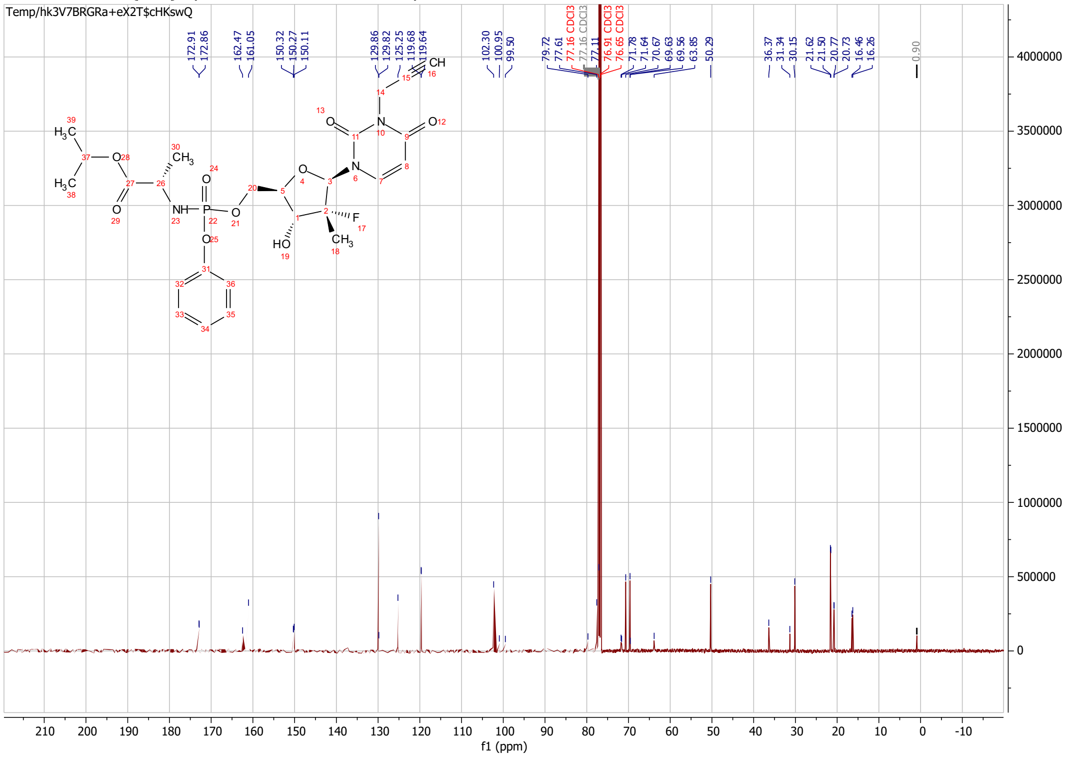

#### Uncropped Western Blot

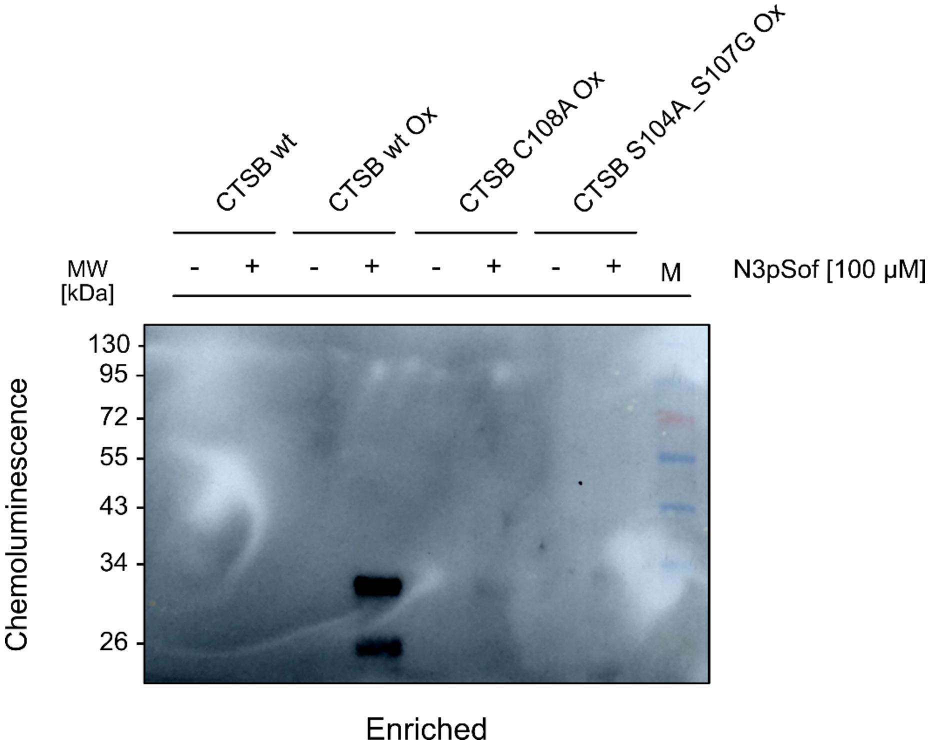
